# Starvation-induced autophagy occurs independently of the ATG1 complex in Chlamydomonas

**DOI:** 10.64898/2026.03.23.713624

**Authors:** Yong Zou, Yixin Wu, Simon Stael, Panagiotis N. Moschou, Xiaohong Zhuang, Elena A. Minina, Peter V. Bozhkov

## Abstract

The survival of eukaryotes during starvation depends on effective nutrient recycling *via* autophagy. Accordingly, loss of autophagy-related (ATG) proteins, including the nutrient-sensing ATG1 kinase complex, typically results in reduced fitness or lethality under nutrient limitation. The green alga *Chlamydomonas reinhardtii* provides a tractable model for autophagy studies, as its ATG repertoire is encoded by single-copy genes. Here, we generated a comprehensive library of *ATG* deletion mutants and examined their growth and autophagy during starvation. Surprisingly, starvation-induced autophagy occurred in the absence of ATG1 complex components (ATG1, ATG11, ATG13, and ATG101), revealing ATG1-independent autophagy and challenging the canonical model for autophagy initiation.

## Background

Autophagy is a major catabolic pathway contributing to intracellular recycling in eukaryotes, with pivotal role in cellular homeostasis, stress adaptation, and development [1, 2]. The formation of a double-membrane vesicle, termed an autophagosome, that sequesters cargo requires the activity of four core molecular modules: the ATG1 initiation complex, the phosphatidylinositol 3-kinase (PI3K) nucleation complex, the ATG9 cycling complex, and two ubiquitin-like ATG8 and ATG12 conjugation systems [3, 4] (Fig. 1a). Although autophagy is well characterized in budding yeast and metazoa, it remains comparatively less understood in photosynthetic organisms, particularly algae, hindering our understanding of the evolution of autophagic mechanisms and functions.

**Fig. 1.**
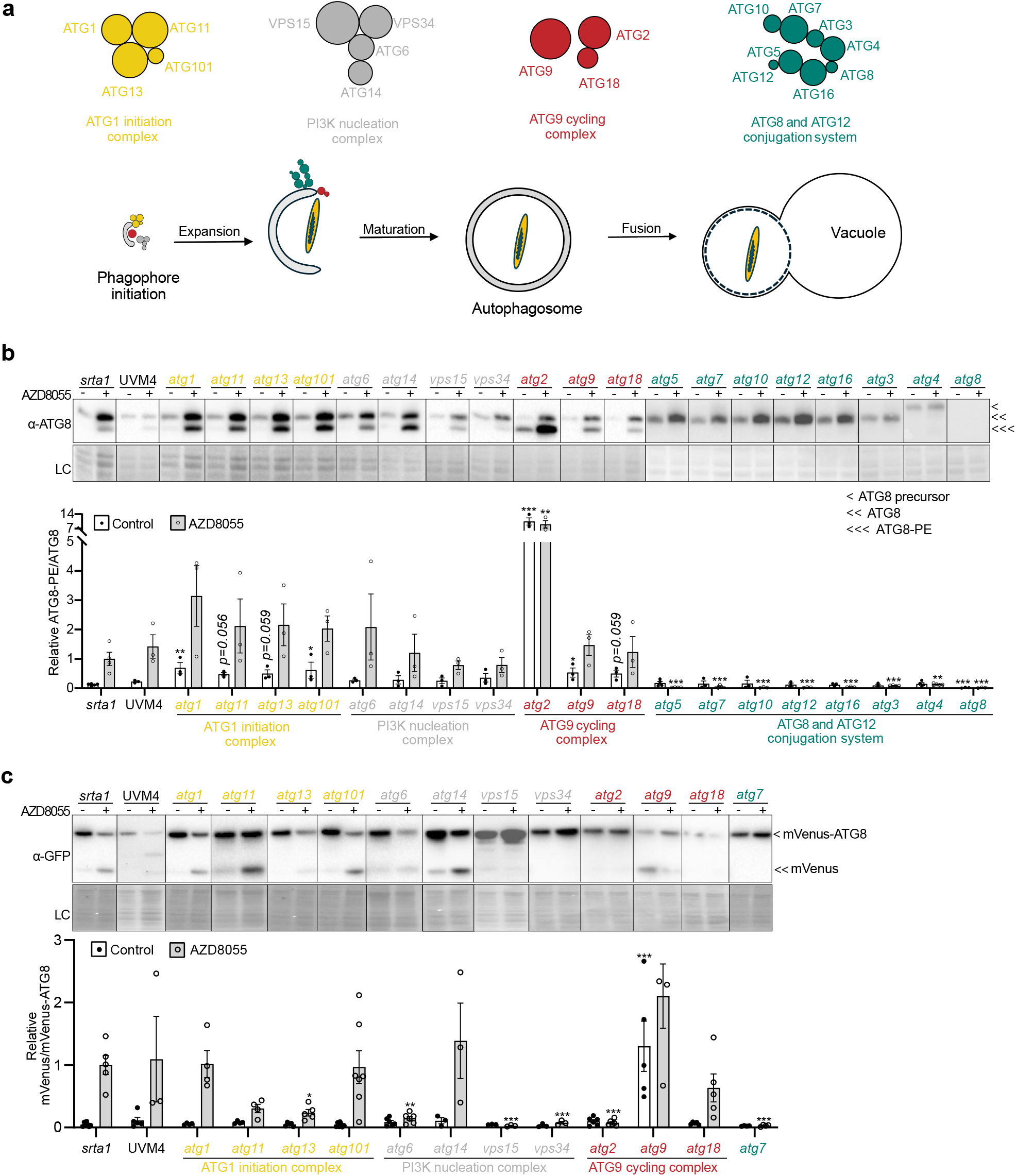
The ATG1 complex is dispensable for starvation-induced autophagy in *C. reinhardtii*. **a**. Schematic representation of the four core molecular modules of the canonical autophagy pathway. **b**. ATG8 lipidation assay of protein extracts from *srta1*, UVM4, and *ATG* deletion strains grown for 24 h with (+) or without (-) 1 µM AZD8055. LC, loading control. The chart shows the abundance of the lipidated form of ATG8 (ATG8-PE) relative to unconjugated ATG8 (ATG8, or ATG8 precursor in the case of *atg4*), quantified by densitometry and normalized to the AZD8055-treated *srta1* strain. **c**. Autophagic flux analysis of *srta1* and *ATG* deletion strains grown for 24 h with (+) or without (-) 1 µM AZD8055 using the mVenus cleavage assay. LC, loading control. The chart shows the abundance of free mVenus relative to mVenus-ATG8 in the corresponding protein extracts, quantified by densitometry and normalized to the AZD8055-treated *srta1* strain. Data in **b** and **c** represent means ± SEM of more than three biological replicates. Autophagy mutants were compared with *srta1* or UVM4 (for *vps15*) using two-way ANOVA followed by Dunnett’s test. *, *p*<0.05; **, *p*<0.01; ***, *p*<0.001. The original, uncropped immunoblot images shown in **b** and **c** are provided in Additional File 1: Fig. S4.

In contrast to land plants (embryophytes), which possess orthologues of all yeast and metazoan core *ATG* genes, currently available genome assemblies of the two green algal clades, charophytes (a clade within streptophytes) and chlorophytes, lack one or more canonical *ATG* genes (Additional file 1: Fig. S1; Additional file 2: Table S1;) [5]. Remarkably, the chlorophyte model alga *Chlamydomonas reinhardtii* retains a complete set of *ATG* genes, making it a valuable system for functional comparative studies across kingdoms and between chlorophyte and streptophyte lineages, including *Arabidopsis thaliana*, the most thoroughly studied plant model for autophagy. Furthermore, the presence of all *ATG* genes as single copies in the *C. reinhardtii* genome facilitates targeted gene ablation.

Here, we conducted a comprehensive functional analysis of autophagy in *C. reinhardtii* by systematically deleting core *ATG* genes and characterizing the autophagic flux, cellular ultrastructure and resulting mutants for viability under nutrient starvation conditions known to induce autophagy in this microalga [6-8]. Our analysis shows that components of the ATG8 and ATG12 conjugation systems are essential for autophagy and cell viability, whereas the requirement for the PI3K nucleation and ATG9 cycling complexes is gene specific. Unexpectedly, deletion of *ATG1, ATG11, ATG13*, or *ATG101* did not impair autophagic activity or algal growth, suggesting that autophagy in *C. reinhardtii* can operate independently of the canonical ATG1 initiation pathway.

## Results and discussion

We determined the complete set of *C. reinhardtii* core *ATG* genes using sequence similarity searches and phylogenetic analyses (Additional file 1: Fig. S1) and implemented CRISPR–Cas9 to establish a comprehensive library of *ATG* knockouts in this species. All mutants, except for *vps15*, were established in strain CC-4533 *srta*, optimized for transgene expression and hereafter referred to as *srta1* (Additional file 1: Fig. S2, S3). The *vps15* knockout had been previously established in the UVM4 strain (Additional file 1: Fig. S3).

Next, we systematically examined the effect of each knockout on autophagic activity using two complementary biochemical assays: detection of ATG8 lipidation and GFP–ATG8 cleavage [9]. ATG8 lipidation — the conjugation of ATG8 to phosphatidylethanolamine (PE) that yields the ATG8–PE adduct — marks initiation of autophagosome formation but alone does not indicate completion of autophagic flux. This is instead revealed by GFP–ATG8 cleavage, which reports cargo degradation in the lytic vacuole [9]. For these assays, autophagy was acutely induced in *C. reinhardtii* cells using AZD8055 (1 µM for 24 h), an inhibitor of Target of Rapamycin (TOR), which canonically induces autophagy through relieving inhibitory phosphorylation of ATG1 complex components [10] and robustly activates autophagy across eukaryotes [11].

The ATG8 lipidation assay demonstrated the expected increase in ATG8–PE accumulation upon AZD8055 treatment in *srta1* and UVM4, whereas ATG8-PE was undetectable in all knockouts of the ATG8 and ATG12 conjugation systems responsible for its synthesis (Fig. 1b, Additional file 1: Fig. S4a) [12-14]. To our surprise, efficient ATG8 lipidation was detected in *atg1, atg11, atg13* and *atg101* revealing that initiation of AZD8055–induced autophagy in *C. reinhardtii* can occur independently of the canonical ATG1 initiation complex (Fig. 1b, Additional file 1: Fig. S4a).

Similarly, the PI3K complex, which is known to recruit the ATG8 lipidation machinery to the growing autophagosome [15], was dispensable for initiation of autophagy under these conditions, as loss of its components did not affect ATG8–PE accumulation upon treatment (Fig.1b, Additional file 1: Fig. S4a). Furthermore, loss of ATG9 cycling complex components had no effect on ATG8–PE accumulation, except for *atg2* (Fig. 1b, Additional file 1: Fig. S4a). Like the ATG1 and PI3K complexes, the ATG9 cycling complex acts upstream of ATG8 lipidation and promotes phagophore membrane expansion through three biochemically distinct and sequential activities. ATG9, a lipid scramblase, anchors the complex at the phagophore rim and recruits the lipid transferase ATG2, which transfers phospholipids from the ER to the growing membrane. ATG2 in turn recruits ATG18, a phosphatidylinositol 3-phosphate (PI3P)-binding protein that coordinates complex assembly with PI3K complex activity and allosterically activates the ATG8 lipidation machinery. Consistent with observations in *A. thaliana* [13], loss of ATG2 resulted in markedly elevated levels of ATG8–PE, suggesting severe mismanagement of membrane resources required for completion of ATG8–PE–decorated autophagosomes in this mutant. In contrast, loss of *ATG9* or *ATG18* had no detectable effect on ATG8–PE levels.

It has not escaped our notice that mutants of the ATG1 initiation and ATG9 cycling complexes exhibited elevated ATG8–PE levels also under control conditions, suggesting either increased basal autophagy or a partial impairment/retardation of autophagic flux (Fig. 1b, Additional file 1: Fig. S4a). To discern which was the case and determine whether the observed autophagy initiation proceeded to completion of flux, we employed GFP-ATG8 cleavage assays, implemented here using an mVenus–ATG8 reporter more suitable for spectral unmixing from autofluorescence in *C. reinhardtii* [16] (Fig. 1c, Additional file 1: Fig. S4b).

Upon AZD8055 treatment, free mVenus accumulated robustly in the *srta1* and UVM4 strains and was undetectable in mutants of ATG8 and ATG12 conjugation systems, confirming assay functionality (Fig. 1c, Additional file 1: Fig. S4b). Remarkably, free mVenus was detectable in knockout lines of all ATG1 initiation complex components, most prominently in *atg1* itself, demonstrating that AZD8055–induced autophagy in *C. reinhardtii* can proceed to completion independently of canonical ATG1 kinase activity. In eukaryotes, AZD8055 is known to trigger autophagy through TOR-mediated phosphorylation of ATG1 complex components [11]. Our observation uncovered that *C. reinhardtii* possesses ATG1-independent mechanisms for transducing the TOR signal.

Notably, mVenus accumulation was significantly reduced in *atg13* and *atg11* mutants compared to *atg1* (Fig. 1c, Additional file 1: Fig. S4b), suggesting that these components may play a role beyond ATG1 complex. No detectable increase in free mVenus under control conditions in mutants of the ATG1 initiation complex confirmed that the elevated ATG8–PE levels observed in these lines (Fig. 1b) result from partial retardation of autophagic flux rather than increased autophagy induction.

Noteworthy, while autophagy initiation was intact in PI3K complex mutants, depletion of any of the PI3K complex components, except ATG14, blocked autophagic flux (Fig.1c; Additional file 1: Fig. S4b). This indicates that ATG1-independent autophagy of *C. reinhardtii* still requires PI3K complex function, consistent with the observations in other eukaryotes [13, 17, 18].

Intriguingly, loss of ATG9 cycling complex components produced a spectrum of effects on autophagic flux; where *atg2* showed abrogated autophagic activity, *atg18* had a slightly reduced flux, and loss of ATG9 resulted in a strong increase in autophagic flux, detectable even under non-inducing conditions (Fig.1c; Additional file 1: Fig. S4b). These results indicate that although *C. reinhardtii* can induce autophagy independently of the ATG1 initiation complex and ATG9, these core ATG proteins are required for maintenance of autophagic homeostasis, as their loss leads to hyperactivation at distinct stages of the pathway. Specifically, loss of the ATG1 initiation complex significantly increased autophagy initiation while maintaining wild-type-like flux, whereas loss of ATG9 drastically elevated autophagic flux with only minor increase in the initial ATG8–PE synthesis.

To confirm that mVenus–ATG8 cleavage in the mutant lines occurred in the lytic vacuole, as inferred from the GFP-cleavage assay, we next employed confocal laser scanning microscopy to examine mVenus localization in the established lines. mVenus remains fluorescent under both the neutral pH of the cytoplasm and the acidic pH of the lytic vacuole, allowing tracking of mVenus–ATG8 translocation through the sequential stages of autophagy and identifying which steps are disrupted in each ATG knockout mutant (Additional file 1: Fig. S5). Under control conditions, lytic vacuoles in *C. reinhardtii* are relatively small and not discernable without additional staining [19] (Fig. 2a). Upon AZD8055 treatment, they become markedly enlarged in an autophagy-independent manner, as evidenced by the vacuolar morphology in *atg7* mutants, and are clearly visible even by bright-field microscopy, facilitating identification of the compartment to which mVenus–ATG8 should be delivered by autophagy (arrows and asterisks, Fig. 2a, Additional file 1: Fig. S6).

**Fig. 2.**
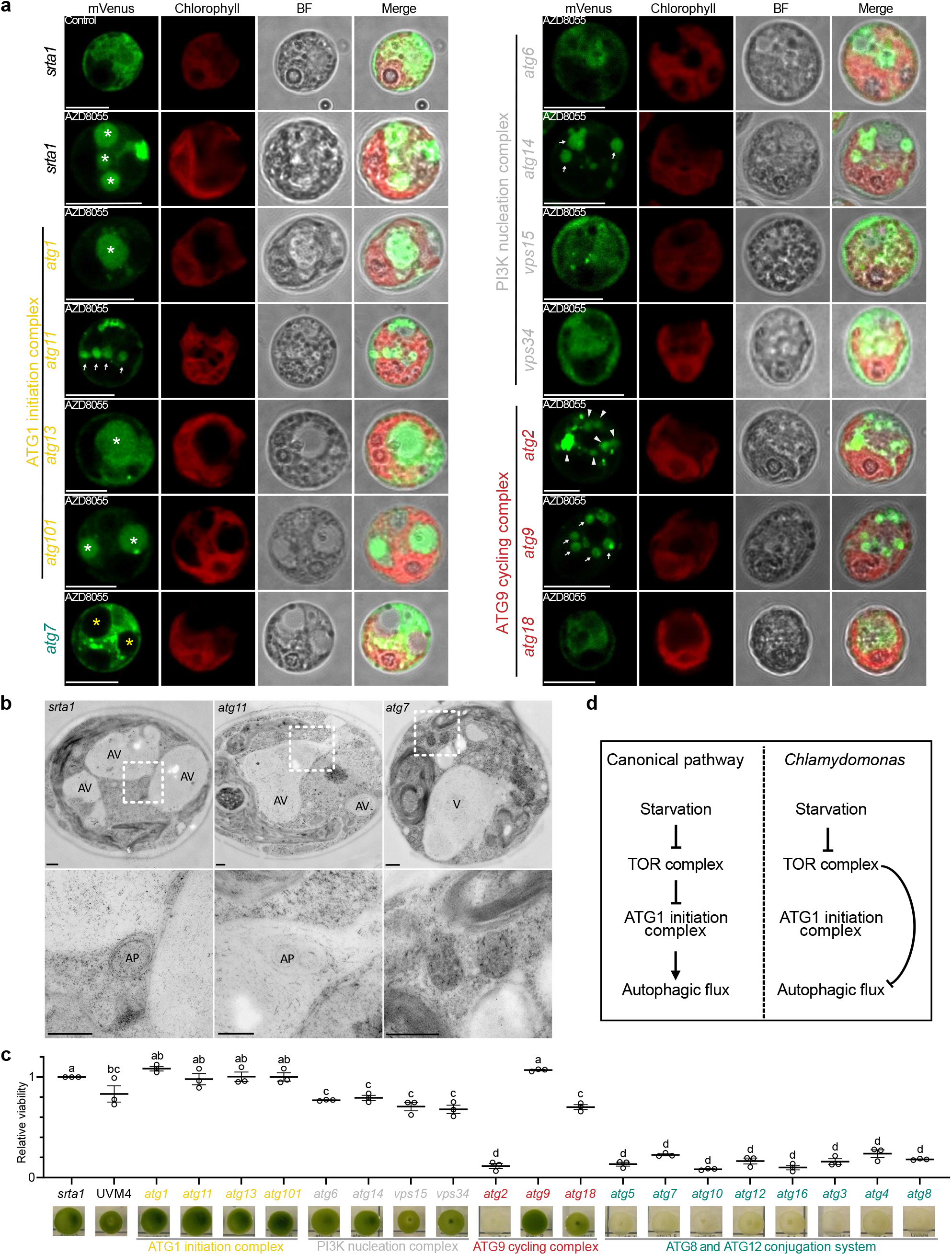
The ATG1 complex is dispensable for autophagosome delivery to the vacuole and for viability during gradual nutrient depletion in *C. reinhardtii*. **a**. Confocal microscopy of mVenus-ATG8 expressing cells shows accumulation of vacuolar mVenus signal (white asterisks or arrows) in response to 5 µM AZD8055 treatment for 2 h. Vacuoles lacking mVenus signal in *atg7* mutant are indicated by yellow asterisks. Abnormal mVenus accumulation in *atg2* mutant is indicated by white triangles. Scale bars, 10 µm. **b**. Transmission electron microscopy reveals autophagosome-like structures in the cytoplasm and vacuoles of the *srta1* and *atg11* strains, but not in the *atg7* strain, after 24 h of treatment with 1 µM AZD8055. Enlarged boxed areas are shown on the right. AP, autophagosome-like structure; AV, autophagic vacuole; V, vacuole. Scale bars, 500 nm. **c**. Representative images of algal colonies of the *srta1*, UVM4, and *ATG* deletion strains after 60 days of continuous cultivation on TAP plates without subculturing (late stationary phase). The chart shows relative viability of the indicated strains, normalized to *srta1*, estimated by quantifying green pigmentation intensity using image analysis. Data represent means ± SEM of three independent measurements. Different letters indicate statistically significant differences among strains; one-way ANOVA followed by Tukey’s honestly significant difference test, *p*<0.05. Colony images from earlier time points during the 60-day cultivation period are provided in Additional File 1: Fig. S8. **d**. Schematic illustration of the major finding that, in contrast to the evolutionarily conserved canonical autophagy pathway, starvation-induced autophagy in *C. reinhardtii* bypasses the ATG1 initiation complex.

As expected, AZD8055 treatment triggered relocation of mVenus from the cytoplasm to the vacuole in *srta1*, but not in *atg7* (Fig. 2a, Additional file 1: Fig. S6). Consistent with the GFP-cleavage assay, mVenus localized to the vacuole in all knockouts of ATG1 complex components, confirming normal autophagic flux, whereas loss of PI3K complex components abolished vacuolar localization, except for *atg14* (Fig. 2a, Additional file 1: Fig. S6).

Interestingly, knockouts of ATG9 cycling complex components exhibited a spectrum of mVenus localization patterns. In *atg2*, the fluorescent signal accumulated on aberrant autophagosomal structures; in *atg18*, it partially translocated to the vacuole; whereas in *atg9*, it was mostly relocated from the cytoplasm to the vacuole (Fig. 2a, Additional file 1: Fig. S6). These observations corroborate the GFP-cleavage and ATG8 lipidation assays and indicate that components of this complex in *C. reinhardtii* perform functions beyond their canonical role in ATG9 cycling, potentially enabling alternative autophagy pathways when individual components are lost.

Additionally, we performed ultrastructural analysis of *C. reinhardtii* cells treated with AZD8055 using transmission electron microscopy. The results confirmed delivery of autophagosome-like structures to the vacuole in the absence of the ATG1 initiation complex (Fig. 2b). Additionally, we observed putative aberrant autophagic structures in the cytoplasm of mutants lacking PI3K complex components or ATG2 (Additional file 1: Fig. S7). Notably, *atg9* and *atg18* mutants displayed distinct vacuolar morphologies: *atg9* showed numerous moderately enlarged vacuoles, whereas *atg18* exhibited vacuolar engulfment of cytoplasmic material (Additional file 1: Fig. S7), suggesting possible activation of different microautophagy routes in these backgrounds.

We next asked whether the ATG1-independent response observed during acute AZD8055-induced autophagy also occurs during gradual nutrient depletion representing physiologically relevant conditions. To this end, we assessed the viability of ATG mutants during prolonged incubation on solidified medium by monitoring signs of early senescence — a characteristic autophagy deficiency phenotype [9, 20]. Expectedly, loss of ATG8 and ATG12 conjugation systems components caused severe early senescence and reduced viability, while background strains retained dark green coloration indicative of active photosynthesis even after 60 days (Fig. 2c; Additional file 1: Fig. S8). This demonstrated that ATG8– PE synthesis is indispensable for autophagy in *C. reinhardtii* under acute and prolonged stress, consistent with its established role across eukaryotes [12-14].

Intriguingly, mutants of the canonical ATG1 initiation complex, including double knockouts, exhibited no detectable acceleration of senescence relative to *srta1* (Fig. 2c; Additional file 1: Fig. S8 and Fig. S9). This contrasts sharply with *A. thaliana*, where the ATG1 complex is critical for normal senescence [21, 22]. To test whether this dispensability was condition-specific, we assessed viability under nitrogen-limited conditions — a stress that also requires ATG1 complex activity in *A. thaliana* [21, 22]. ATG1 initiation complex mutants again showed wild-type-like viability in *C. reinhardtii* (Additional file 1: Fig. S10). Together, these results indicate that the dispensability of the ATG1 initiation complex in *C. reinhardtii* is not limited to acute stress or standard growth conditions, pointing to a fundamental difference in autophagy regulation compared to vascular plants (Fig. 2d).

Consistently with the observations during AZD8055-induced autophagy, loss of ATG2 caused severe early senescence, whereas loss of ATG9 and ATG18 produced wild-type-like and intermediate phenotypes respectively, revealing strikingly unequal contributions of the ATG9 cycling complex components (Fig. 2c; Additional file 1: Fig. S8). These results suggest that the ATG8 lipidation machinery can be activated independently of ATG18 and of ATG9-mediated complex recruitment to the phagophore. The severity of the *atg2* phenotype may also partly reflect autophagy-independent roles of ATG2-mediated lipid transfer.

Most strikingly, all mutants of PI3K complex showed only moderate decrease in viability upon prolonged nutrient depletion, in stark contrast to its requirement for autophagic activity under acute stress (Fig. 1c; Additional file 1: Fig. S4b; Fig. 2a; Additional file 1: Fig. S6; Fig. S7), indicating compensatory mechanisms upregulating autophagy under mild prolonged stress.

## Conclusions

Our systematic genetic dissection of the autophagy machinery in *C. reinhardtii* revealed an unexpected versatility of this pathway. Consistent with observations in other eukaryotes, synthesis of ATG8–PE proved critical for autophagic activity under both acute and prolonged mild stress, rendering all components of the ATG8 and ATG12 conjugation systems essential for autophagy. However, under all tested conditions, including acute induction via TOR inhibition, autophagy proceeded independently of the ATG1 complex, challenging the canonical model of ATG1-dependent autophagy initiation.

Similarly, other so-called core ATGs, such as ATG9 and ATG18, appear to be dispensable for autophagy in *C. reinhardtii*. Nevertheless, we hypothesize that these components remain conserved because of their roles in maintaining autophagic homeostasis, as their loss resulted in skewed initiation or completion of autophagic activity.

These findings uncover a previously unrecognized divergence in autophagy initiation within the green lineage, distinguishing *C. reinhardtii* from land plants such as *A. thaliana* and other eukaryotic systems in which the ATG1 complex is indispensable under nutrient-limited conditions. Our results support the existence of an alternative ATG1-independent mechanism initiating autophagy in chlorophytes. Together, this work highlights the evolutionary plasticity of autophagy regulation and establishes *C. reinhardtii* as a powerful model system for dissecting non-canonical autophagy pathways in a genetically streamlined photosynthetic eukaryote.

## Methods

### Genome survey and phylogenetic analysis of ATG proteins

*A. thaliana* ATG protein sequences were used as queries (corresponding gene accession number in Additional file 2: Table S2) for BLAST searches against selected genomes in Phytozome [23] or UniProt [24]. Putative orthologues were identified using reciprocal BLAST searches. Protein sequences were aligned in MEGA X [25], and phylogenetic trees shown in Additional file 1: Fig. S1 were constructed using the neighbor-joining method with default parameters. A total of 1,000 bootstrap replicates were performed, and branch support values greater than 50% are shown. Gene structures presented in Additional file 1: Fig. 3a and 3b were visualized using GSDS 2.0 [26].

### Background strains and growth conditions

CC-4533, its derivative *srta1*, and UVM4 were used as genetic backgrounds for genome editing. *C. reinhardtii* cells were cultured under standard condition: axenically in Tris-Acetate-Phosphate (TAP) liquid or solid medium at 22 °C and under a 16 h light/8 h dark cycle with a light intensity of 110 μmol m^−2^ s^−1^. Cells in the stationary phase were diluted 20-fold into fresh medium and grown for one day, followed by treatment with 1 µM AZD8055 (Santa Cruz Biotechnology, sc-364424). For growth under nitrogen-limited conditions, TAP-0.75N medium containing 25% of the nitrogen content of full-strength TAP was used.

### *ATG* gene deletion by CRISPR-Cas9

A CRISPR-Cas9 based targeted insertional mutagenesis approach [27] was used to knock out genes of interest. For each gene, a single guide RNA (sgRNA, Additional file 2: Table S3) was designed to target one of the first four exons. Cells were electroporated with the ribonucleoprotein complex and a donor DNA fragment containing an expression cassette. The cassette included the Hsp70/RBCS2 synthetic promoter, a selectable marker (*ble* for zeocin, *AphVII* for hygromycin, or *AphVIII* for paromomycin), and the RBCS2 terminator. Transformants were screened by colony PCR (Additional file 1: Fig. S3, primers listed in Additional file 2: Table S3). All strains generated in this study are listed in Additional file 2: Table S4.

### mVenus-ATG8 expression

Constructs used for expressing mVenus-ATG8 were generated using the MoClo kit [28]. L1 vectors were assembled from the following L0 modules: the *PsaD* promoter (pCM0-016), *mVenus* gene (pCM0-066), the genomic sequence of *CrATG8* [20], and the *PsaD* terminator (pCM0-114). L2 vectors carrying hygromycin or paromomycin resistance were subsequently assembled. The L2 vectors were introduced into WT or *ATG* deletion strains using the glass bead method [29]. Transformants were screened by confocal microscopy and immunoblot analysis. Plasmids are listed in Additional file 2: Table S5.

### Growth assays

Cells from plates were pre-cultured to the stationary phase. Equal numbers of cells (2.5 × 10^5^) from each strain were loaded onto plates for growth assays. Plates were maintained in the growth chamber under the above-described conditions, and the growth phenotypes were photographed on designated days. Viability assays were performed as previously described [20]. Briefly, RGB images of cell growth after 60 days were separated into red, green, and blue channels, and the green channel was used for subsequent analysis. Circular regions of interest (ROIs) containing cells were defined to quantify color intensity. Background intensity was determined as the mean value of four randomly selected regions lacking cells. For each strain, the corrected color intensity was calculated by subtracting the background value from the measured intensity. Finally, color intensity values were normalized to the average value of the background strain.

### ATG8 lipidation assay

One milliliter of cell cultures in log phase (2–10 × 10^6^ cells mL^−1^) was harvested by centrifugation at 17,000 × g for 1 min at room temperature and snap-frozen in liquid nitrogen. The cell pellet was resuspended in 50 μL 2 × SDS sample buffer [30] and incubated at 70 °C for 5 min. Samples were vortexed at maximum speed for 30 s and then centrifuged at 17,000 × g for 15 min at room temperature. The resulting supernatant was diluted with an equal volume of distilled water and used for immunoblotting. Protein crude extracts were separated on manually cast 15% SDS-PAGE gels supplemented with 6 M urea [31] and transferred to Trans-Blot Turbo Midi 0.2 µm PVDF membranes (BioRad, 1704157). Membrane was blocked for 1 h in 5% skimmed milk in PBST buffer. Primary anti-ATG8 (Agrisera, AS14 2769, 1:5,000) was incubated with the membranes overnight, followed by incubation with secondary anti-rabbit antibody (Agrisera, AS09 602, 1:20,000) for 1 h before detection. Densitometric analysis of the western blots was performed using the software Image Lab 6.1 (BioRad).

### Autophagic flux assay

Crude protein extracts were prepared as described for the ATG8 lipidation assay. Proteins were separated on 4–20% TGX Stain-Free precast gels (Bio-Rad, 4568096). Primary anti-GFP (Roche, 11814460001; 1:2,000) and secondary anti-mouse antibody (Agrisera, AS11 1772; 1:10,000) were applied. Densitometric analysis of the western blots was performed using the software Image Lab 6.1 (BioRad).

### Confocal microscopy

The mVenus fluorescent signal was detected using a confocal microscopy (Leica Stellaris 5) with excitation at 514 nm and emission collected from 540 to 590 nm. Because cells became fragile after 24 h of treatment with 1 µM AZD8055, the treatment duration for confocal imaging was shortened to 2 h, while the concentration of AZD8055 was increased to 5 µM.

### High pressure freezing (HPF) and transmission electron microscopy (TEM)

The general procedures for preparing the TEM samples and ultra-thin sectioning of samples were conducted as previously described [32, 33], with some modifications. In brief, two-day *C. reinhardtii* cells were centrifuged at 2,000 g for 5 min, resuspended in 0.15 M mannitol dissolved in TAP, and fixed using a high-pressure freezer (EM ICE, Leica). The frozen samples were then transferred to an AFS2 temperature-controlling system (Leica) for freezing substitution in acetone with 0.25% glutaraldehyde and 0.1% uranyl acetate at −80°C for 24□h. Subsequently, the samples were rinsed with precooled acetone, and slowly warmed up to −45°C followed by embedding in HM20 resin and polymerization under ultraviolet light. Sections were subjected to post-staining with aqueous uranyl acetate/lead citrate and examined under a Hitachi HT-7800 microscope (Hitachi High-Technologies Corporation, Japan).

### Statistical analysis

Statistical analyses were performed as described in figure legends. Raw data are provided in Additional file 2: Table S6.

## Supplementary Information

The online version contains supplementary material available at

Additional file 1. Supplementary figures S1-S10.

Additional file 2. Supplementary tables S1-S6.

## Authors’ contributions

Conceptualization and study design: Y.Z., E.A.M., P.V.B. Supervision: S.S., E.A.M., P.V.B. Funding acquisition: S.S., P.N.M., E.A.M., P.V.B. Experiments and data analysis: Y.Z., Y.W., X.Z. Results interpretation: Y.Z., E.A.M., P.V.B. Manuscript draft: Y.Z., P.N.M., E.A.M., P.V.B. Manuscript review and editing: All authors. All authors read and approved the final manuscript.

## Data availability

All data supporting the findings of this study are available within the paper and its Supplementary Information. If any raw data files are required in a different format, they are available from the corresponding authors upon request.

