## Additional file 1 for "Starvation-induced autophagy occurs independently of the ATG1 complex in Chlamydomonas"

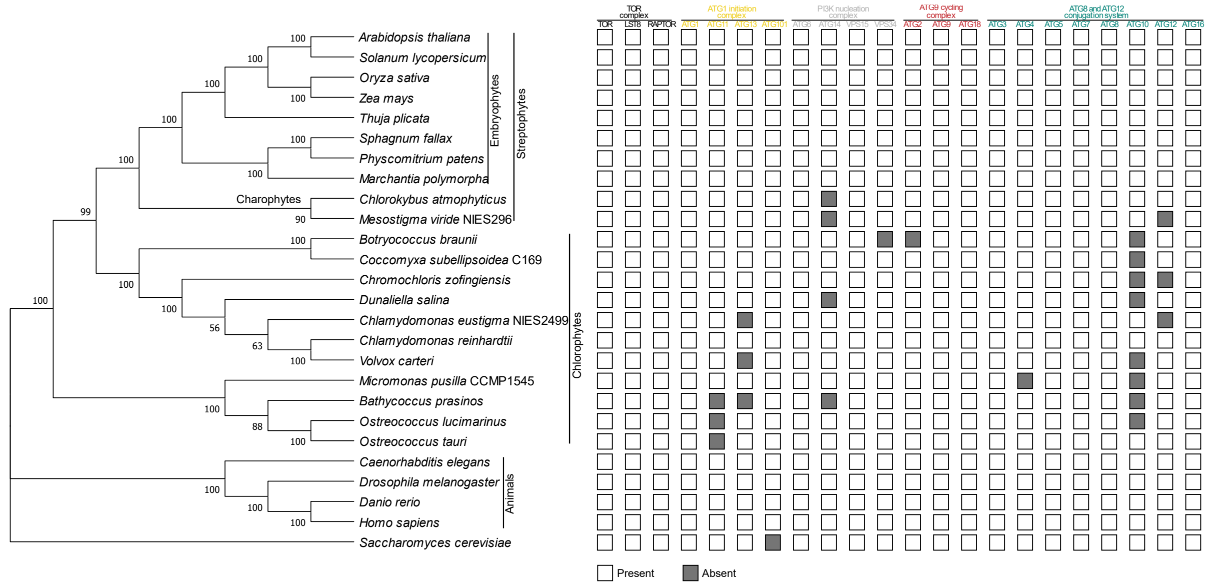


**Fig. S1.** Both green algal clades (Charophytes and Chlorophytes) lack certain core *ATG* genes, with *C. reinhardtii* retaining a complete set.

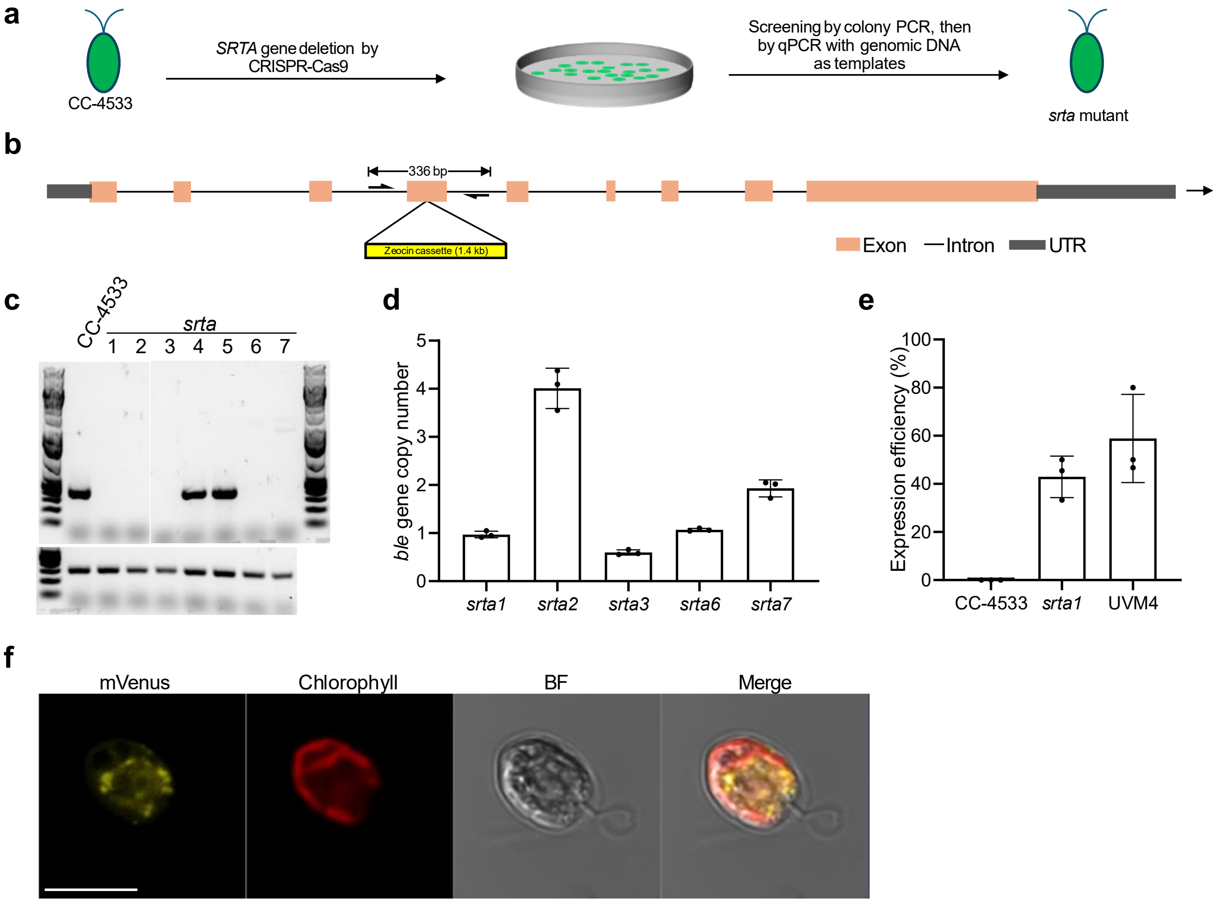


**Fig. S2.** *srta1* background strain is engineered for robust expression of foreign genes.

**a.** Schematic illustration of *srta1* strain generation. CC-4533 was used as the starting strain for genome editing.

**b.** Schematic map of the *SRTA* gene editing site, showing a pair of primers (half arrows) flanking the target region, used for PCR analysis.

**c.** Colony PCR of several potential transformants using the primers shown in **b**. The absence of the 384-bp amplicon present in the CC-4533 strain indicates potential gene deletion event following CRISPR–Cas9-mediated genomic cleavage. The positive technical control below shows successful amplification of the *CrMCA-II* gene [1].

**d.** Real-time quantitative PCR analysis of *ble* (for Zeocin resistance) and *RACK1* genes (reference, set to 1) using genomic DNA as a template to evaluate the number of genome insertion events. Data represent means ± SEM of triplicate measurements. *srta1*, *srta3*, and *srta6* each harbored a single *ble* gene copy, indicating single insertion events, whereas *srta2* and *srta7* appeared to contain two and more insertions.

**e.** Expression efficiency of mVenus in strains CC-4533, *srta1*, and UVM4, quantified as a percentage of transformants exhibiting yellow fluorescence. Data represent means ± SEM of three independent transformation experiments.

**f.** Representative confocal image showing mVenus expression in the *srta1* background strain. BF, bright field. Scale bar, 10 μM.

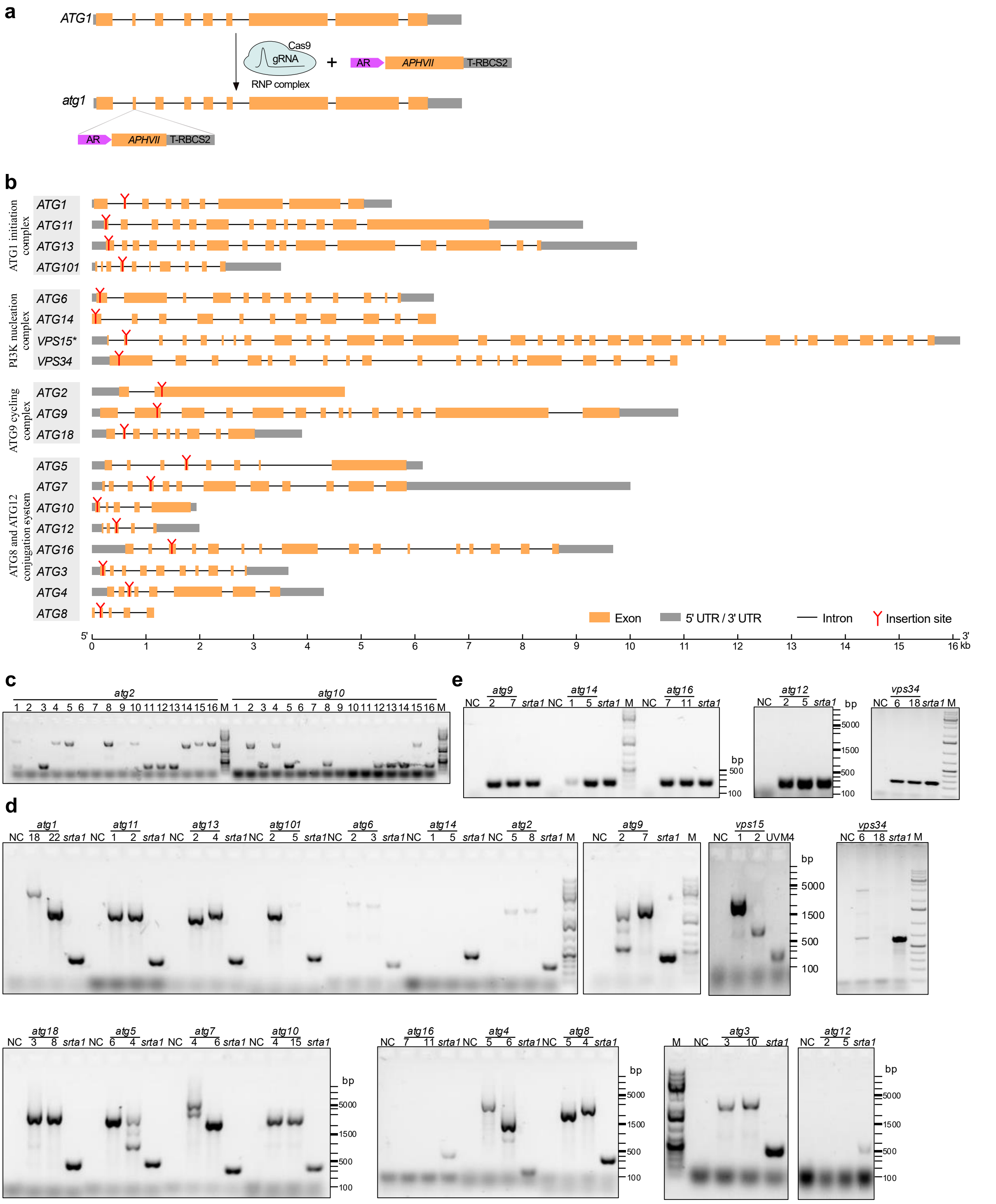


**Fig. S3.** CRISPR–Cas9–mediated deletion of all annotated *ATG* genes in *C. reinhardtii*.

**a.** Schematic illustration of the CRISPR–Cas9 approach used to generate *atg* mutant strains in the *srta1* background.

**b.** Diagram showing the insertion sites of the *AphVII* cassette (resistant to paromomycin) in each targeted *ATG* gene. Insertion sites (indicated by “Y”) were positioned in one of the first four exons of each gene to ensure complete knockout. An asterisk (*) denotes *vps15*, the only mutant generated in the UVM4 background, rather than *srta1*, with *AphVIII* cassette (resistant to hygromycin) insertion.

**c.** Representative results for *atg2* and *atg10* from the initial screening of transformants by colony PCR using crude DNA extracts.
**d.** Confirmation of selected *atg* mutants by PCR using purified genomic DNA as template. Distilled water served as a negative control (NC), and *srta1* or UVM4 (for *vps15*) were used as positive controls. M, DNA marker.
**e.** Verification of PCR-negative *atg* strains through amplification of the *RACK1* gene to rule out poor genomic DNA quality.

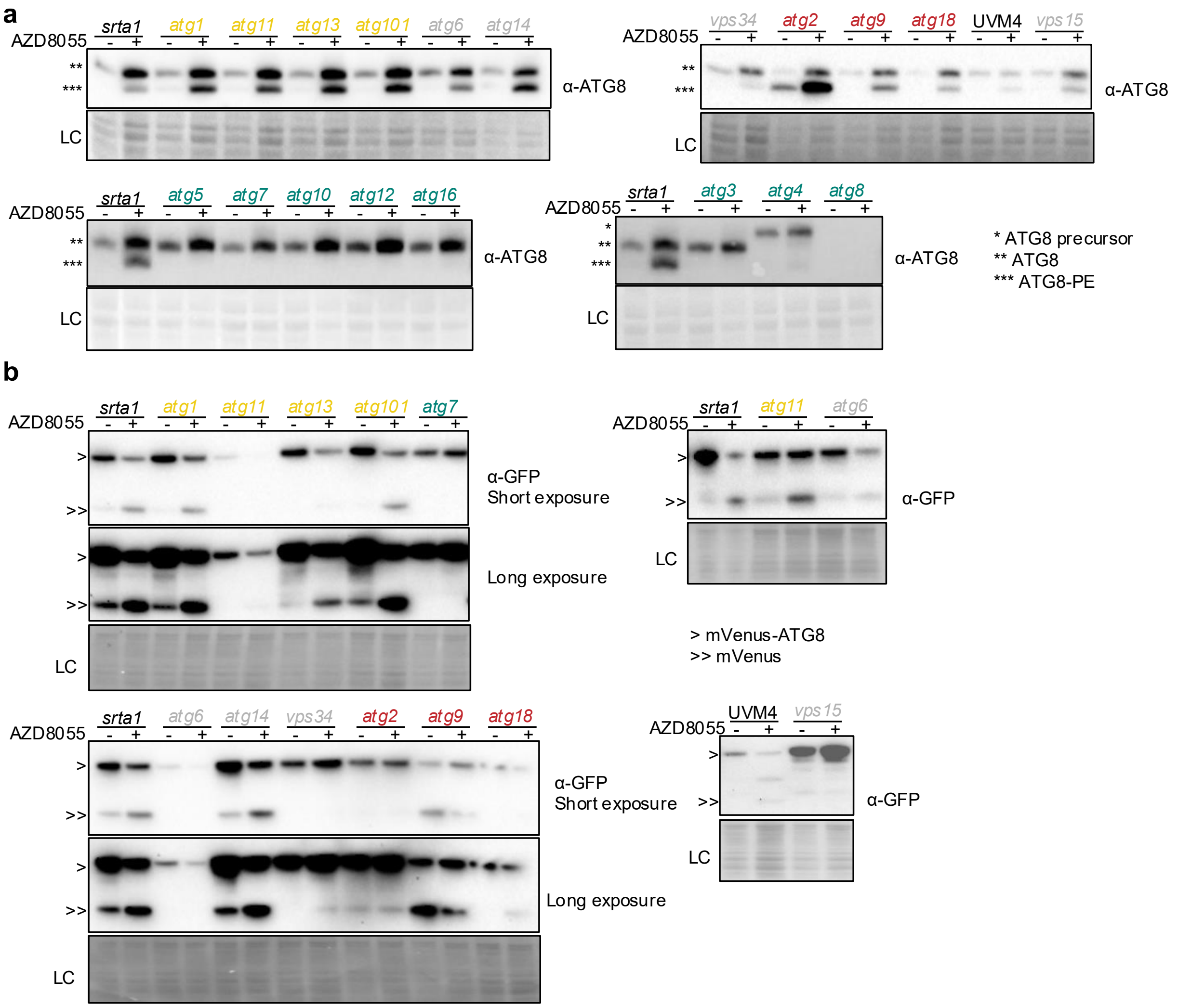


**Fig. S4.** The ATG1 initiation complex is dispensable for starvation-induced autophagy in *C. reinhardtii.* Related to Fig. 1c and d, which show cropped and re-arranged immunoblots.

**a.** ATG8 lipidation assay of protein extracts from *srta1*, UVM4, and ATG deletion strains grown for 24 h with (+) or without (-) 1 µM AZD8055. LC, loading control.
**b.** Autophagic flux analysis of *srta1* and ATG deletion strains grown for 24 h with (+) or without (-) 1 µM AZD8055 using the mVenus cleavage assay. LC, loading control.

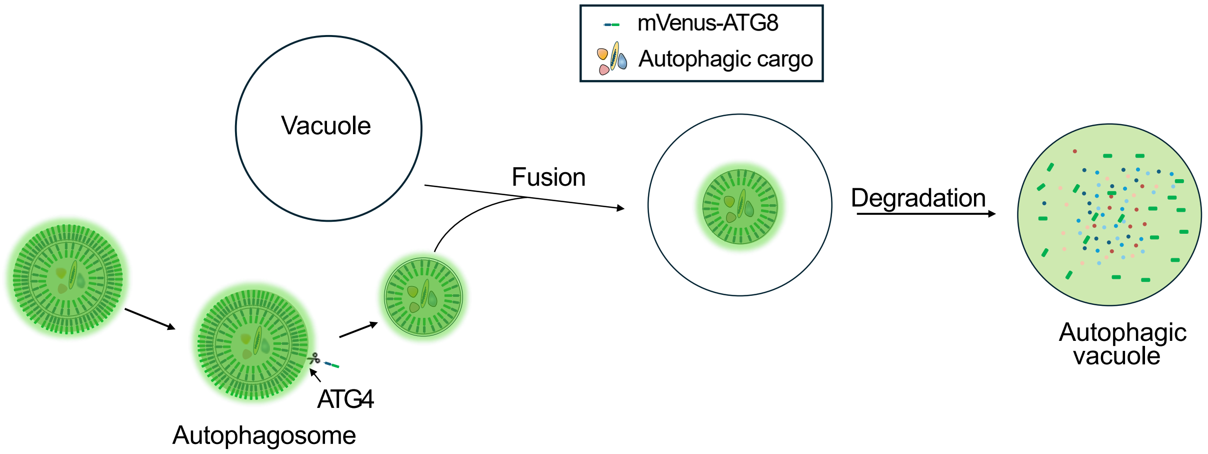


**Fig. S5.** Schematic illustration of autophagic vacuole formation accompanied by vacuolar accumulation of mVenus-Atg8. mVenus–ATG8 labels both the outer and inner membranes of the autophagosome. ATG8 conjugated to the outer membrane is cleaved by ATG4 and recycled. Upon autophagosome–vacuole fusion, the inner membrane and enclosed cargo are delivered into the vacuolar lumen. There, ATG8 is degraded, whereas mVenus is relatively resistant to vacuolar proteolysis. As a result, mVenus accumulates within the vacuole, serving as a marker of autophagic flux in both the mVenus cleavage assay and confocal microscopy. Related to Figs. 1d, 2a, S8b, and S9.

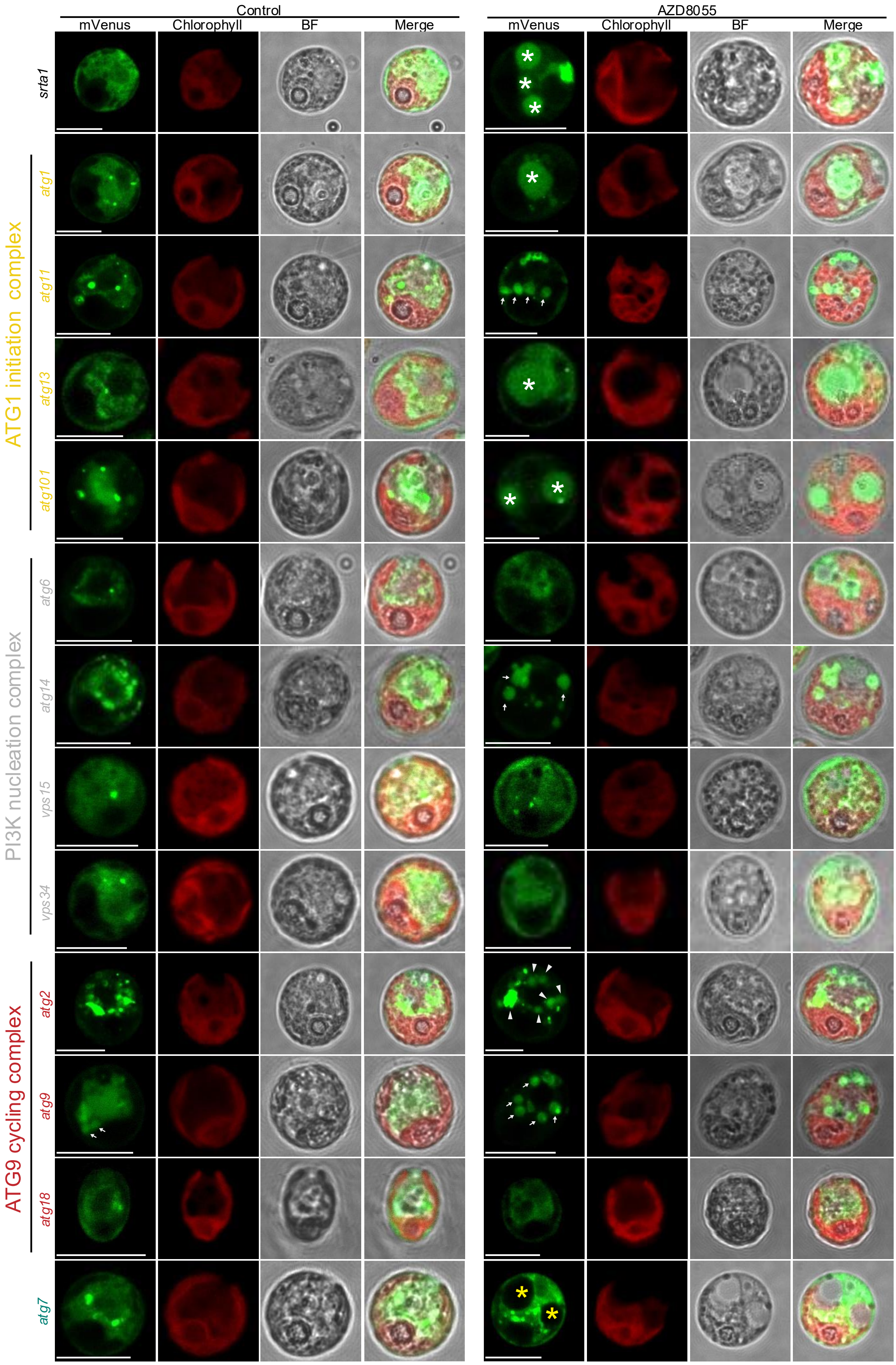


**Fig. S6.** Confocal microscopy of mVenus-ATG8 expressing cells of *srta1* strain and autophagy mutants, treated with the presence or absence of AZD8055 (5 μM

AZD8055 for 2 h). mVenus signal, white asterisks or arrows; vacuoles lacking mVenus signal, yellow asterisks; abnormal mVenus accumulation: white triangle. Scale bars, 10 μm. Related to Fig. 2a.

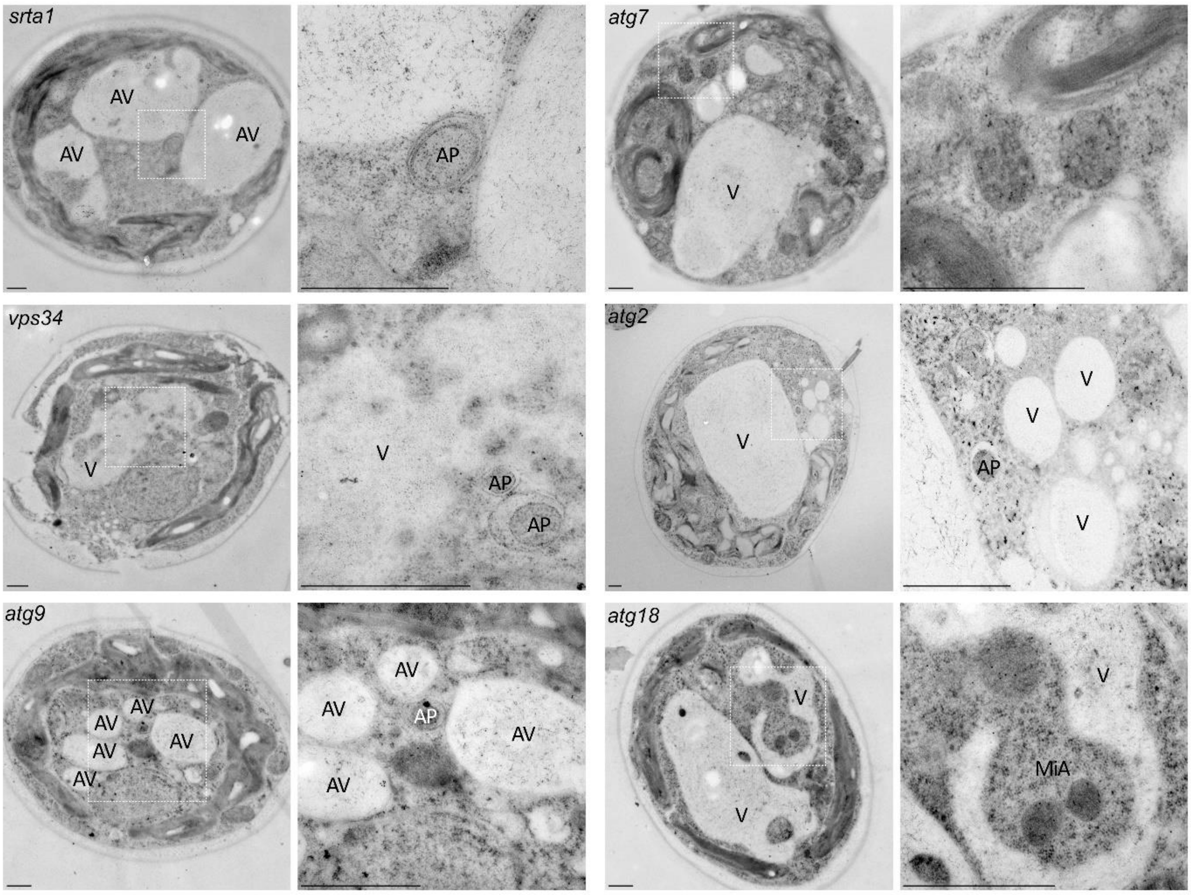


**Fig. S7.** Transmission electron microscopy of *vps34*, *atg2*, *atg9*, and *atg18* strains treated with 1 μM AZD8055 for 24 h reveals autophagosome-like structures in the *vps34*, *atg2* and *atg9* strains, and microautophagy-like event in *atg18* strains. Enlarged boxed areas are shown on the right. AP, autophagosome-like structure; AV, autophagic vacuole; V, vacuole; MiA, microautophagy-like event. Scale bars, 500 nm. Related to Fig. 2b.

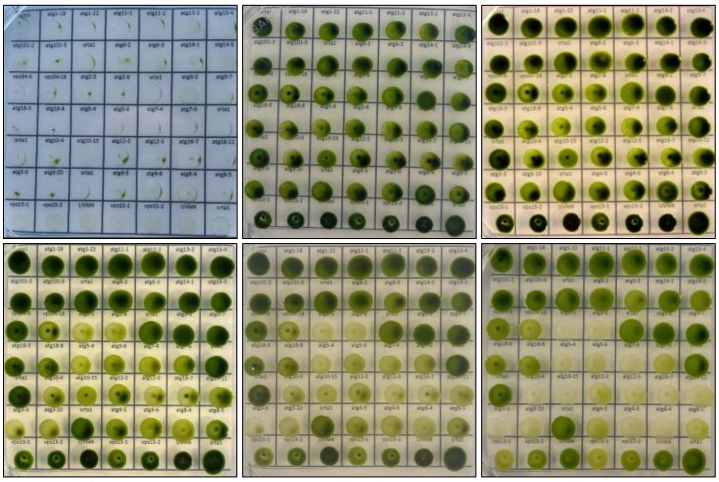


| | | | | | | |
|---|---|---|---|---|---|---|
| *srta1* | *atg1-18* | *atg1-22* | *atg11-1* | *atg11-2* | *atg13-2* | *atg13-4* |
| *atg101-2* | *atg101-5* | *srta1* | *atg6-2* | *atg6-3* | *atg14-1* | *atg14-5* |
| *vps34-6* | *vps34-18* | *atg2-5* | *atg2-8* | *srta1* | *atg9-2* | *atg9-7* |
| *atg18-3* | *atg18-8* | *atg5-4* | *atg5-6* | *atg7-4* | *atg7-6* | *srta1* |
| *srta1* | *atg10-4* | *atg10-15* | *atg12-2* | *atg12-5* | *atg16-7* | *atg16-11* |
| *atg3-3* | *atg3-10* | *srta1* | *atg4-5* | *atg4-6* | *atg8-4* | *atg8-5* |
| *vps15-1* | *vps15-2* | UVM4 | *vps15-1* | *vps15-2* | UVM4 | *srta1* |

**Fig. S8.** The ATG1 initiation complex is dispensable for stationary-phase viability in *C. reinhardtii.* Equal numbers of cells ($2.5 \times 10^5$) of *srta1*, UVM4, and ATG deletion strains were inoculated onto TAP plates (day 0) and maintained without subculturing for 60 days. Photographs were taken at the indicated time points. The inoculation scheme defining the position of individual strains on the plates is shown on the right. Related to Fig. 1b.

**a**

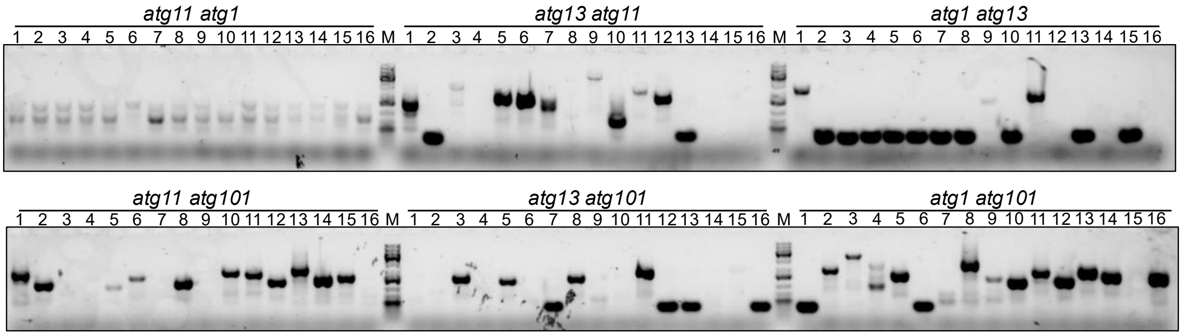


**b**

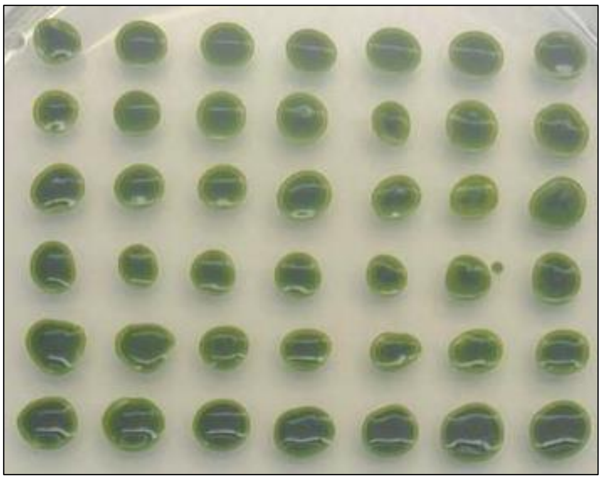

| | | | | | | |
|---|---|---|---|---|---|---|
| *atg11 atg1.1* | *atg11 atg1.2* | *atg11 atg1.3* | *atg11 atg1.4* | *atg11 atg1.5* | *atg11 atg1.6* | *atg11 atg1.7* |
| *atg13 atg11.1* | *atg13 atg11.3* | *atg13 atg11.5* | *atg13 atg11.6* | *atg13 atg11.7* | *atg13 atg11.9* | *atg13 atg11.11* |
| *atg1 atg13.1* | *atg1 atg13.9* | *atg1 atg13.11* | *atg1 atg13.12* | *atg1 atg13.14* | *atg1 atg13.16* | *srta1* |
| *atg11 atg101.1* | *atg11 atg101.2* | *atg11 atg101.5* | *atg11 atg101.6* | *atg11 atg101.8* | *atg11 atg101.10* | *atg11 atg101.11* |
| *atg13 atg101.3* | *atg13 atg101.5* | *atg13 atg101.8* | *atg13 atg101.11* | *atg13 atg101.1* | *atg13 atg101.2* | *atg13 atg101.4* |
| *atg1 atg101.2* | *atg1 atg101.3* | *atg1 atg101.4* | *atg1 atg101.5* | *atg1 atg101.8* | *atg1 atg101.9* | *atg1 atg101.10* |

**Fig. S9.** ATG1 complex double mutants of *C. reinhardtii* exhibit no stationary phase growth phenotype.

**a.** Colony PCR verification of the ATG1 complex double mutants.

**b.** Colonies of ATG1 complex double mutants and *srta1* strain after 40 days of growth on TAP plates.

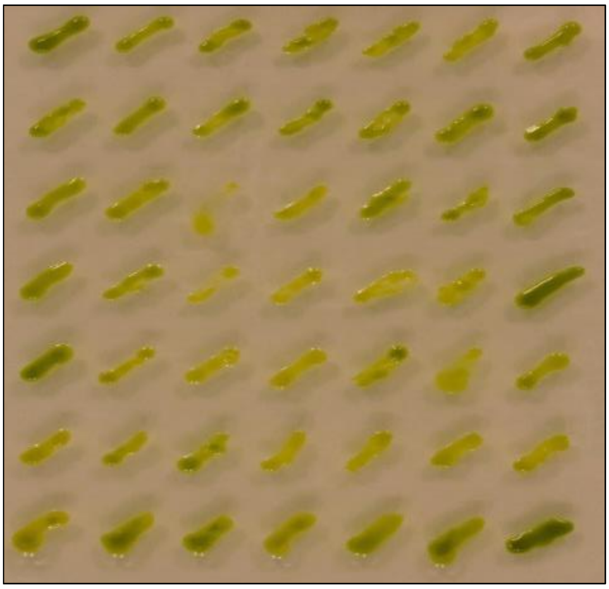

| | | | | | | |
|---|---|---|---|---|---|---|
| *srta1* | *atg1-18* | *atg1-22* | *atg11-1* | *atg11-2* | *atg13-2* | *atg13-4* |
| *atg101-2* | *atg101-5* | *srta1* | *atg6-2* | *atg6-3* | *atg14-1* | *atg14-5* |
| *vps34-6* | *vps34-18* | *atg2-5* | *atg2-8* | *srta1* | *atg9-2* | *atg9-7* |
| *atg18-3* | *atg18-8* | *atg5-4* | *atg5-6* | *atg7-4* | *atg7-6* | *srta1* |
| *srta1* | *atg10-4* | *atg10-15* | *atg12-2* | *atg12-5* | *atg16-7* | *atg16-11* |
| *atg3-3* | *atg3-10* | *srta1* | *atg4-5* | *atg4-6* | *atg8-4* | *atg8-5* |
| *vps15-1* | *vps15-2* | UVM4 | *vps15-1* | *vps15-2* | UVM4 | *srta1* |

**Fig. S10.** ATG1 complex mutants of *C. reinhardtii* exhibit no growth phenotype under nitrogen-limited conditions. *C. reinhardtii* cultures were grown on complete TAP medium for 10 days and then transferred to the nitrogen-depleted TAP medium (25% of standard nitrogen content) for an additional 10 days before imaging.
